# A Distributed Architectural Code Governs DNA Repair

**DOI:** 10.64898/2026.10.16.759706

**Authors:** Anisha Hundal, Defne Urman, Araad Algouneh, Rehna Krishnan, Chris A. Jordan, Sergei Rudnizky, Adam T. Rybczynski, Tamari Chkuaseli, Anne Hakem, Jun Hao, Philipp G. Maass, Taekjip Ha, Razq Hakem, Karim Mekhail

## Abstract

Repair of DNA double-strand breaks (DSBs) depends on large-scale changes in nuclear shape and genome organization. How damage signaling coordinates such remodeling remains unclear. We find that the human DNA damage response (DDR) couples interdependent, temporally ordered tubulin modifications to nuclear membrane and lipid regulation, driving nuclear reshaping (nucleomorphosis) through DSB-capturing nuclear envelope tubules (dsbNETs). Tubulin detyrosination by vasohibins 1/2 (VASH1/2) and acetylation by α-tubulin acetyltransferase 1 (ATAT1) promote dsbNET formation and support both homologous recombination and non-homologous end joining. Microtubule-dependent processes also promote the phosphorylation-dependent inactivation of the membrane stress-sensing lipid regulator CTP cytidylyltransferase α (CCTα), which typically restrains baseline nuclear lipid droplets (nLDs) and dsbNETs while supporting ATR signaling. Damage-induced nLD formation requires diacylglycerol acyltransferases 1/2 (DGAT1/2) and supports dsbNETs. Our findings reveal a distributed architectural code spanning interconnected cytoskeletal, membrane, and lipid regulators. This code governs nucleomorphosis and its role in DNA repair and genome stability.

## INTRODUCTION

Genome maintenance depends on DNA repair enzymes that recognize and process lesions within chromatin^1–3^. Yet efficient repair also requires cytoskeletal and nuclear envelope components and involves large-scale changes in nuclear shape and genome organization^3^. How damage signaling coordinates repair reactions at lesions with such extensive architectural remodeling remains poorly understood.

Molecular codes provide a means of translating combinations of modifications into biological outcomes. The histone code^4–6^, RNA polymerase II C-terminal domain code^7–10^, and tubulin code^11–16^ exemplify how regulatory information can arise from relationships among modifications, their temporal organization, and their interpretation by dedicated factors. Deciphering such a code requires identifying its elements and determining how their individual functions and interactions generate coordinated outputs.

Whether the DNA damage response (DDR) uses analogous coding elements distributed across broad cellular structures to mediate repair via large-scale nuclear and genome reshaping remains unknown. More specifically, the key unresolved questions are whether diverse architectural modifications exist and crosstalk, how their activities promote or restrain nuclear remodeling, and whether their contributions to DNA repair outcomes are functionally distinct.

We recently identified DNA double-strand break (DSB)-capturing nuclear envelope tubules (dsbNETs), which promote genome maintenance through reversible, damage-induced nuclear reshaping, or nucleomorphosis^3,17^. Here, using dsbNETs as an experimental entry point, we identified a distributed cytoskeletal–nuclear membrane architectural code. This code couples interdependent, temporally ordered tubulin modifications with the phosphoregulation of a membrane curvature stress and lipid regulator and with the formation of nuclear lipid droplets. Code components cross-regulate and provide non-identical contributions to homologous recombination and non-homologous end joining. Our findings reveal a distributed, functionally integrated architectural code governing the induction, repair functions, and restraint of nucleomorphosis to preserve genome stability.

## RESULTS

### DNA damage triggers a coordinated architectural code

To determine whether DNA damage induces coordinated architectural responses, we profiled the temporal responses of multiple cytoskeletal and membrane remodeling processes following genotoxic stress. Tubulin post-translational modifications regulate distinct microtubule properties^12^. In particular, tubulin detyrosination modulates interactions with various microtubule-binding proteins^18–22^, tubulin acetylation increases microtubules’ mechanical resilience^23,24^, and tubulin polyglutamylation recruits diverse proteins to microtubules and can mediate their graded severing^16,25,26^. We therefore monitored acetylated (Ac-Tub)^14,23,24^, detyrosinated (deTyr-Tub)^13,18,19^, and polyglutamylated (polyE-Tub)^16,25,27^ tubulin following etoposide treatment^17^. Etoposide induced robust and transient increases in deTyr-Tub and Ac-Tub, whereas polyE-Tub displayed a distinct temporal profile, with a modest qualitative increase followed by a significant decline (**Figures 1A–1B and S1A–S1C**). We detected increases in deTyr-Tub and Ac-Tub levels within 5-15 min and 10-15 min of etoposide treatment, respectively (**Figure 1A**). The increases and subsequent decreases in deTyr-Tub and Ac-Tub broadly paralleled γH2AX accumulation and clearance as well as dsbNET formation and reversal (**Figures 1A–1D and S1A–S1B**). The induction of tubulin modifications also showed a dose-dependent response reflecting the magnitude of DNA damage induced (**Figure S1C**). DNA damage induced similar tubulin changes in different cell types (**Figure S1D**). These changes are consistent with DNA damage inducing several tubulin states in a manner consistent with the microtubule-dependent mechanical reshaping of the nuclear envelope underlying dsbNET-driven DSB repair^17^.

**Figure 1.**
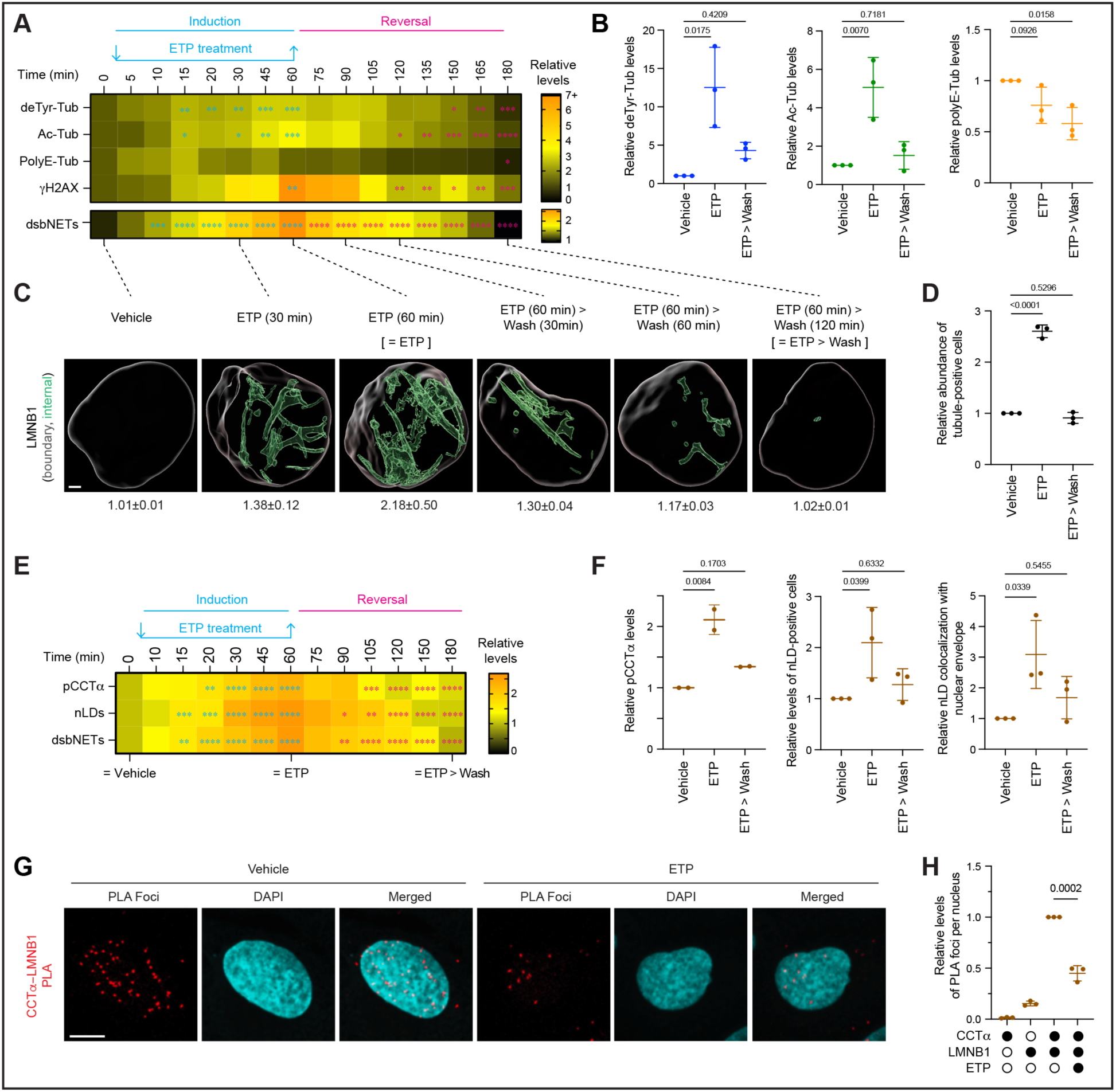
DNA damage reconfigures cytoskeletal and nuclear architectural elements. (A) Heatmap of tubule-positive cells, tubulin PTMs (detyrosination (deTyr-Tub), acetylation (Ac-Tub), polyglutamylation (polyE-Tub)), and γH2Ax levels during etoposide (ETP) treatment and recovery. (B) Quantification of tubulin PTMs under the indicated treatment conditions. (C) Representative 3D reconstructions of LMNB1-marked nuclear architectures and corresponding tubule scores (mean ± SEM). n = 5 nuclei per timepoint. (D) Relative levels of tubule-positive cells. (E) Heatmap of tubule-positive cells, phosphorylated CCTα (pCCTα), and nuclear lipid droplet (nLD) kinetics. (F) Quantification of pCCTα, nLD frequency and abundance, and nLD colocalization with the nuclear envelope. (G, H) Representative proximity ligation assay (PLA) images (G) and quantification (H) of CCTα–LMNB1 interactions ± ETP, using antibodies (filled circles) for both proteins together and separately as controls. (A-H) Experiments were performed in U2OS cells treated with 100 µM ETP for 1 h unless otherwise indicated; for wash conditions, cells recovered for 2 h post-ETP unless otherwise specified; Immunoblot signals were normalized to α-tubulin or total CCTα, where applicable; Data are mean ± SD and n = 3 biological replicates unless otherwise indicated; Two-way ANOVA with Tukey’s (B, D, F) or Dunnett’s (A, C) multiple comparisons test; Cyan and magenta asterisks indicate comparisons with 0 min and 60 min, respectively (A, C); ns, P > 0.05; *P ≤ 0.05; **P ≤ 0.01; ***P ≤ 0.001; ****P ≤ 0.0001. Scale bar 2 µm (C) and 10 µm (G).

We next examined the membrane phospholipid regulator CTP:phosphocholine cytidylyltransferase α (CCTα), which couples sensing of membrane lipid composition and curvature stress to phosphatidylcholine synthesis^28^. We also examined nuclear lipid droplets (nLDs)^29^, which may supply lipids necessary for nucleomorphosis. CCTα and nLDs also crosstalk; nLDs can recruit and locally activate CCTα^30,31^ while CCTα-dependent phosphatidylcholine synthesis stabilizes lipid-droplet surface monolayers and limits the formation of enlarged droplets^32^. Of note, CCTα membrane association can be limited by multisite phosphorylation^31,33^, but its contribution to damage-induced nuclear remodeling remains unknown. We observed that DNA damage increased CCTα phosphorylation (pCCTα) as detected using an antibody raised against phosphorylated S315 and S319 (**Figures 1E–1F and S1E**). DNA damage also decreased CCTα proximity to the nuclear periphery marker LMNB1 (**Figures 1G–1H**), consistent with phosphorylation-associated release of CCTα from the nuclear envelope^28,33^. pCCTα induction coincided with nLD accumulation and increasing nLD colocalization with the nuclear envelope at the periphery or dsbNETs (**Figures 1E-1F and S1F-S1G**). These observations are consistent with a model in which CCTα inactivation permits nLD accumulation, creating a source of lipid precursors for nuclear membrane expansion during nucleomorphosis.

Similar changes in tubulin modifications, pCCTα, and nLDs occurred when DNA damage was induced with zeocin or ionizing radiation (**Figures S1H-S1P**). The transient increases in pCCTα and nLDs also broadly paralleled γH2AX accumulation and clearance and dsbNET formation and reversal during DNA repair (**Figures 1A–1F and S1A–S1P**). DNA damage also increased cytosolic phospholipase A2 (cPLA2) localization at the nuclear periphery (**Figure S1Q**), consistent with increased nuclear envelope lipid remodeling^34^. Unlike the addition of exogenous oleate, which promotes CCTα recruitment to the nuclear envelope by large-scale changes in membrane electrostatic charge^31^, DNA damage did not induce comparable envelope enrichment of CCTα (**Figure S1R**).

Our results reveal a temporally organized response spanning tubulin modifications, CCTα phosphoregulation, and nLD dynamics, suggesting a distributed architectural code during nucleomorphosis. The observed structural states tracked DNA damage and repair progression, while the magnitude of their induction correlated with genotoxic dose and γH2AX accumulation. The tubulin- and membrane-related changes observed are consistent with increased microtubule-dependent nuclear envelope reshaping and suggest CCTα inactivation alters nuclear envelope characteristics while boosting nLD levels providing a source of lipid precursors for nuclear membrane expansion during nucleomorphosis. Collectively, our findings suggest the existence of a coordinated architectural code that is dynamically rewritten during the DNA damage response.

### Distinct architectural code elements specify distinct repair outcomes

A defining feature of biological codes is that distinct code elements produce distinct outputs. We therefore studied how individual cytoskeletal and membrane elements impact DSB repair. Depletion of α-tubulin acetyltransferase-1 (ATAT1, which deposit Ac-Tub)^14,23,24^ or the tubulin tyrosine carboxypeptidases vasohibin-1 or -2 (VASH1/2, which promote deTyr-Tub)^13,18,19^ impaired both HR and NHEJ repair (**Figures 2A, 2B, and S2A**). ATAT1 depletion decreases AcTub, abrogates dsbNET induction, and disrupts the efficiency and kinetics of DSB repair^17^. Consistent with a broader reconfiguration of tubulin states in DSB repair, depletion of VASH1/2 also abrogated dsbNET induction (**Figure 2C**) and increased γH2AX accumulation (**Figure 2D**). Depletion of ATAT1, VASH1, or VASH2 increased DSB burden and delayed repair kinetics, as measured by neutral comet tail moments (**Figures S2B and S2C**) and impaired the resection-indicating RPA2 phosphorylation overall and in EdU-positive cells (**Figures S2D and S2E**). These phenotypes occurred despite small changes in cell-cycle distribution (**Figure S2F**) and did not reflect reduced damage induction, as control and depleted cells exhibited similarly rapid initial chromosome fragmentation following etoposide exposure (**Figure S2G**). Pharmacological inhibition of VASH1/2 with EPOY^18^ also suppressed etoposide-induced dsbNET formation and increased γH2AX foci (**Figures S2A, S2H and S2I**). Together, our observations are consistent with VASH1/2-dependent deTyr-Tub and ATAT1-dependent Ac-Tub broadly supporting both major DSB repair pathways.

**Figure 2.**
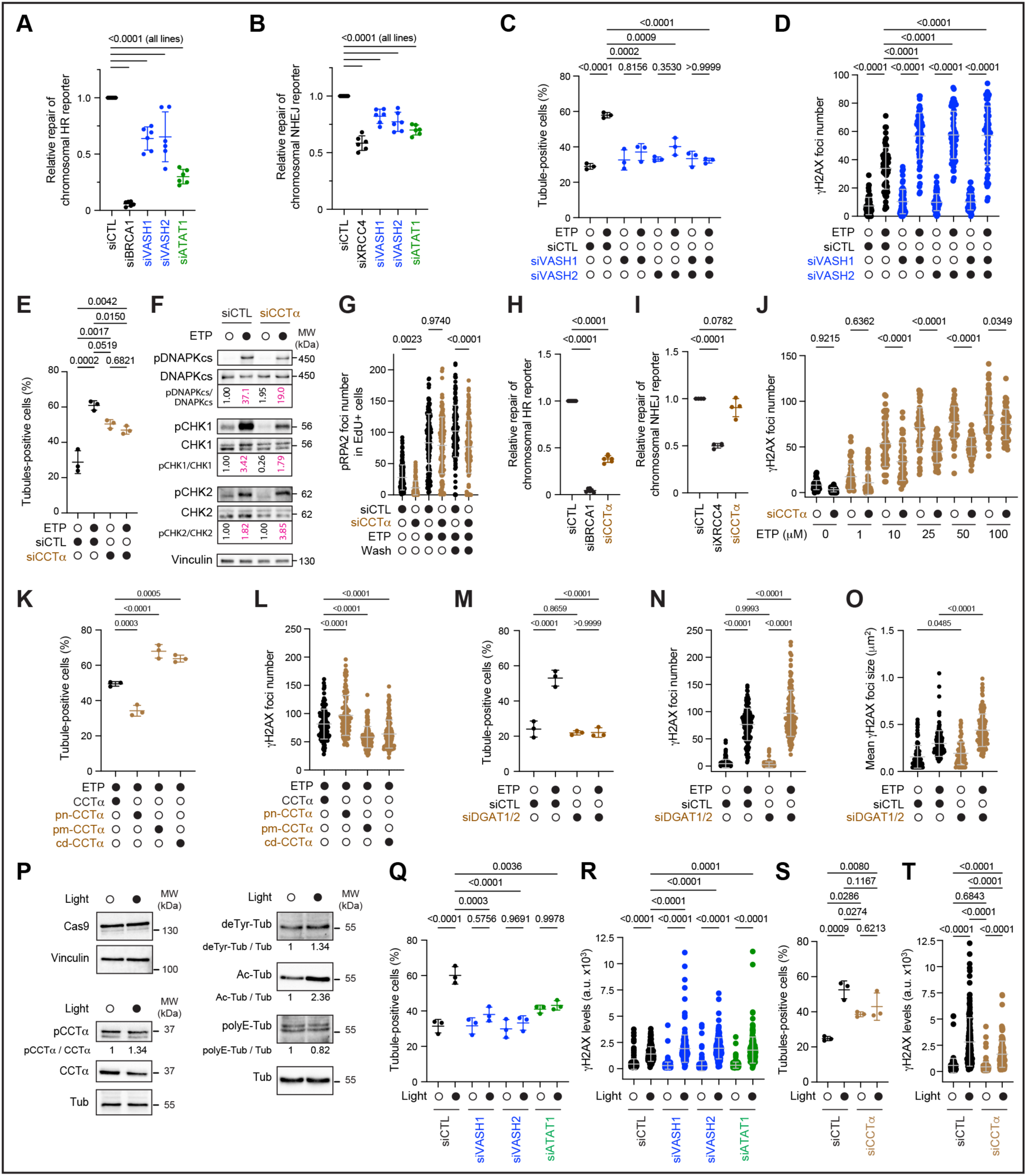
Diverse impacts of architectural changes on DNA repair. (A,B) HR (A) and NHEJ (B) efficiencies following VASH1, VASH2, or ATAT1 depletion, measured using DR-GFP and EJ5 reporters, respectively. BRCA1 and XRCC4 depletion served as controls. (C, D) Percentage of tubule-positive cells (C) and γH2AX foci number (D) following VASH1 and/or VASH2 depletion ± ETP. (E) Percentage of tubule-positive cells following CCTα depletion in ± ETP. (F) Immunoblot of DDR kinase substrate phosphorylation following CCTα depletion ± ETP. (G) Phospho-RPA2 (Ser4/Ser6) foci number in EdU-labeled cells following CCTα depletion ± ETP and 4h post-ETP recovery (Wash). (H, I) HR (H) and NHEJ (I) efficiencies following CCTα depletion. (J) γH2Ax foci number following CCTα depletion and treatment with the indicated ETP concentrations. (K, L) Percentage of tubule-positive cells (K) and γH2AX foci number (L) following endogenous CCTα depletion and re-expression of wild-type CCTα, phosphonull-CCTα (pn-CCTα), phosphomimetic-CCTα (pm-CCTα) or catalytic-dead CCTα (cd-CCTα) followed by ETP treatment. (M–O) Percentage of tubule-positive cells (M), γH2AX foci number (N), and foci size (O) following combined DGAT1/DGAT2 depletion ± ETP. (P) Immunoblot analysis of Cas9, pCCTα, and tubulin PTMs following UV-activated AluGG cgRNA-mediated DNA damage. (Q–T) Percentage of nuclear envelope tubule-positive cells (Q, S) and γH2AX nuclear intensity (R, T) following VASH1, VASH2, or ATAT1 depletion (Q, R), or CCTα depletion (S, T), with or without UV illumination (Light). (A-T) Data are mean ± SD; n = 6 (A, B), 3 (C–E, G–O, Q–T), 2 (F), or 1 (P) biological replicates; One-way ANOVA with Dunnett’s (A, B, H, I) or two-way ANOVA with Tukey’s (C–E, G, J–O, Q–T) multiple comparisons test; Identical Cas9/cgRNA delivery and UV illumination conditions were used (P–T).

CCTα depletion produced a strikingly different phenotype. Loss of CCTα hyper-induced dsbNETs in the absence of exogenous damage but prevented their further induction following etoposide treatment (**Figure 2E**), as would be expected upon depleting a negative regulator of dsbNET formation. This phenotype was specific to the nuclear CCTα as depletion of its CCTβ non-nuclear isoform^33^ did not alter dsbNET formation (**Figure S2J**). Thus, rather than simply blocking nucleomorphosis, CCTα depletion shifts the nuclear envelope into a state permissive to elevated baseline dsbNETs, blunting its standard damage-dependent induction.

Because mechanical tension can activate DDR kinases at the nuclear envelope^3,35,36^ and our data suggested that CCTα knockdown remodels the nuclear envelope even before exposure to DNA damage, we asked whether DDR kinase activation is altered in CCTα-depleted cells. Indeed, CCTα depletion reduced damage-dependent activation of ATR/CHK1 and DNA-PKcs and concordantly decreased RPA2 phosphorylation, whereas ATM/CHK2 activation was comparatively preserved (**Figures 2F, 2G, and S2K**). Functionally, CCTα depletion appeared to impair HR without significantly altering NHEJ (**Figures 2H and 2I**), distinguishing this repair phenotype from the combined HR and NHEJ defects caused by disrupting ATAT1 or VASH1/2 (**Figures 2A and 2B**). CCTα-depleted cells also displayed a partial shift from S to G1 phase (**Figure S2L**), consistent with the reduced engagement of HR.

We next examined the consequences of this altered repair response for DSB persistence. Depleting CCTα decreased damage-induced γH2AX foci without altering clustered-foci-indicating focus size across different etoposide concentrations (**Figures 2J and S2M**) and reduced neutral comet tail moments (**Figures S2N and S2O**), consistent with decreased DNA damage levels. The observed reductions did not reflect decreased damage induction, as control and CCTα-depleted cells displayed similarly rapid initial chromosome fragmentation following shorter exposure to etoposide (**Figure S2G**). Instead, detectable DSB-associated signals resolved more rapidly following CCTα depletion (**Figure S2O**). We observed similar results following ionizing radiation or zeocin treatment, as CCTα depletion increased baseline dsbNETs, prevented their further induction following genotoxic stress, and lowered DNA damage signals (**Figures S2P–S2S**). Together, our results indicate that CCTα depletion lowers overall DSB burden. This phenotype occurs despite similar damage induction and reduced reporter DSB repair by HR but not by NHEJ, suggesting the induction of NHEJ or other similarly fast-acting repair mechanisms, either broadly or in a more targeted manner across the genome.

Next, to assess the functional significance of CCTα phosphorylation, we first depleted endogenous CCTα using a 3′UTR-targeting siRNA. We then expressed EGFP-tagged wild-type CCTα, catalytically inactive CCTα (cd-CCTα with the K122R mutation)^37^, or phosphonull or phosphomimetic CCTα (pn-CCTα and pm-CCTα with all membrane binding--related phosphosites [S315/S319/S321/S322/S323] modified to A or D, respectively)^31,33^. pn-CCTα reduced dsbNET abundance relative to wild-type CCTα in etoposide-treated cells, whereas pm-CCTα and cd-CCTα increased dsbNET formation (**Figure 2K**). This supports our model that the catalytic activity of CCTα is important for curbing unscheduled dsbNET formation. In etoposide-treated cells, pm-CCTα and cd-CCTα significantly reduced γH2AX accumulation, which was instead boosted by pn-CCTα (**Figure 2L**). Our results indicate that CCTα phosphostate, which dictates catalytic activity, differentially controls nucleomorphosis and γH2AX accumulation, with phosphomimetic and catalytically inactive CCTα supporting dsbNET formation and markedly limiting damage-induced γH2AX, recapitulating the phenotype observed in CCTα knockdown conditions.

The depletion or phosphorylation-dependent inactivation of CCTα may increase dsbNET formation by rendering the nuclear membrane more malleable and/or by increasing the abundance of nLDs providing phospholipids to growing dsbNETs. Indeed, the membrane-related processes impacting nucleomorphosis were not only sensitive to direct CCTα modulation. Specifically, depletion of the nLD biogenesis factors diacylglycerol O-acyltransferase 1 and 2 (DGAT1/2)^38^ suppressed damage-induced dsbNET formation and increased γH2AX foci number (**Figures 2M–2O**) without altering initial damage induction or cell-cycle distribution (**Figures S2G and S2L**). Pharmacological inhibition of fatty-acyl CoA synthetase (ACSL), an additional lipid droplet biogenesis factor upstream of the DGAT1/2 proteins^39^, with Triacsin-C similarly prevented etoposide-dependent dsbNET induction (**Figure S2T**). Thus, the membrane-remodeling layer encompasses the dsbNET-restraining CCTα and the dsbNET-promoting nLDs, which are themselves promoted by DGAT1/2 but restrained by CCTα. Therefore, disruption of different elements of the membrane-regulatory layer produces different architectural and repair consequences.

Finally, we asked whether the observed relationships persisted when DSBs were generated independently of chemical or radiation-induced genotoxic stress. Indeed, light-activated very-fast CRISPR (vfCRISPR) targeting Alu elements^40,41^ rapidly induced similar tubulin modifications and pCCTα changes (**Figure 2P**). Depleting ATAT1, VASH1, or VASH2 again suppressed dsbNET induction and exacerbated γH2AX accumulation following vfCRISPR activation (**Figures 2Q and 2R**). In contrast, as expected, CCTα depletion increased basal dsbNETs, prevented their further induction following vfCRISPR activation, and decreased subsequent γH2AX accumulation (**Figures 2S and 2T**). Thus, the consequences of perturbing key cytoskeletal and membrane code elements were recapitulated upon acute, targeted DSB induction and are not limited to pharmacologically induced genotoxic stress.

Collectively, our findings indicate that disrupting distinct architectural elements yields different repair outcomes. Whereas loss of ATAT1 or VASH1/2 broadly impaired HR and NHEJ and promoted persistent DNA damage, loss of CCTα altered basal nucleomorphosis, attenuated damage-induced DDR signaling, and appeared to alter the relative reliance on different repair pathways, culminating in the accelerated disappearance of detectable DSBs. Our data indicate that cytoskeletal and membrane elements are not functionally interchangeable but instead influence the upstream signaling, efficiency, and utilization of DSB repair pathways. Thus, the elements identified herein on different components of the cellular architecture function as parts of a code that serves an instructive role for nucleomorphosis and DNA repair.

### DNA damage response kinases establish the architectural code

Having established that distinct architectural code elements differentially contribute to DNA repair, we next examined how DNA damage regulates these processes^1,17^. Notably, inhibition of ATM or ATR, but not DNA-PKcs, impaired damage-induced deTyr-Tub accumulation (**Figures 3A and 3B**). Microtubule-associated proteins (MAPs) can regulate VASH-mediated detyrosination by controlling VASH access to the microtubule surface, including under stress conditions such as ischemia^12,42,43^. Also, publicly available datasets identified DNA damage-dependent phosphorylation of MAP1B (**Figure S3A**)^44^. We therefore examined MAP1B and VASH1 association with tubulin. DNA damage decreased MAP1B– tubulin proximity signals while increasing VASH1–tubulin signals (**Figures 3C and 3D**). Moreover, the damage-induced decrease in MAP1B–tubulin proximity required ATM and ATR, but not DNA-PKcs activity (**Figure 3E**). The results are consistent with ATM/ATR signaling, but not DNA-PKcs signaling, promoting a transition from MAP1B-associated to VASH-remodeled microtubules during the DNA damage response.

**Figure 3.**
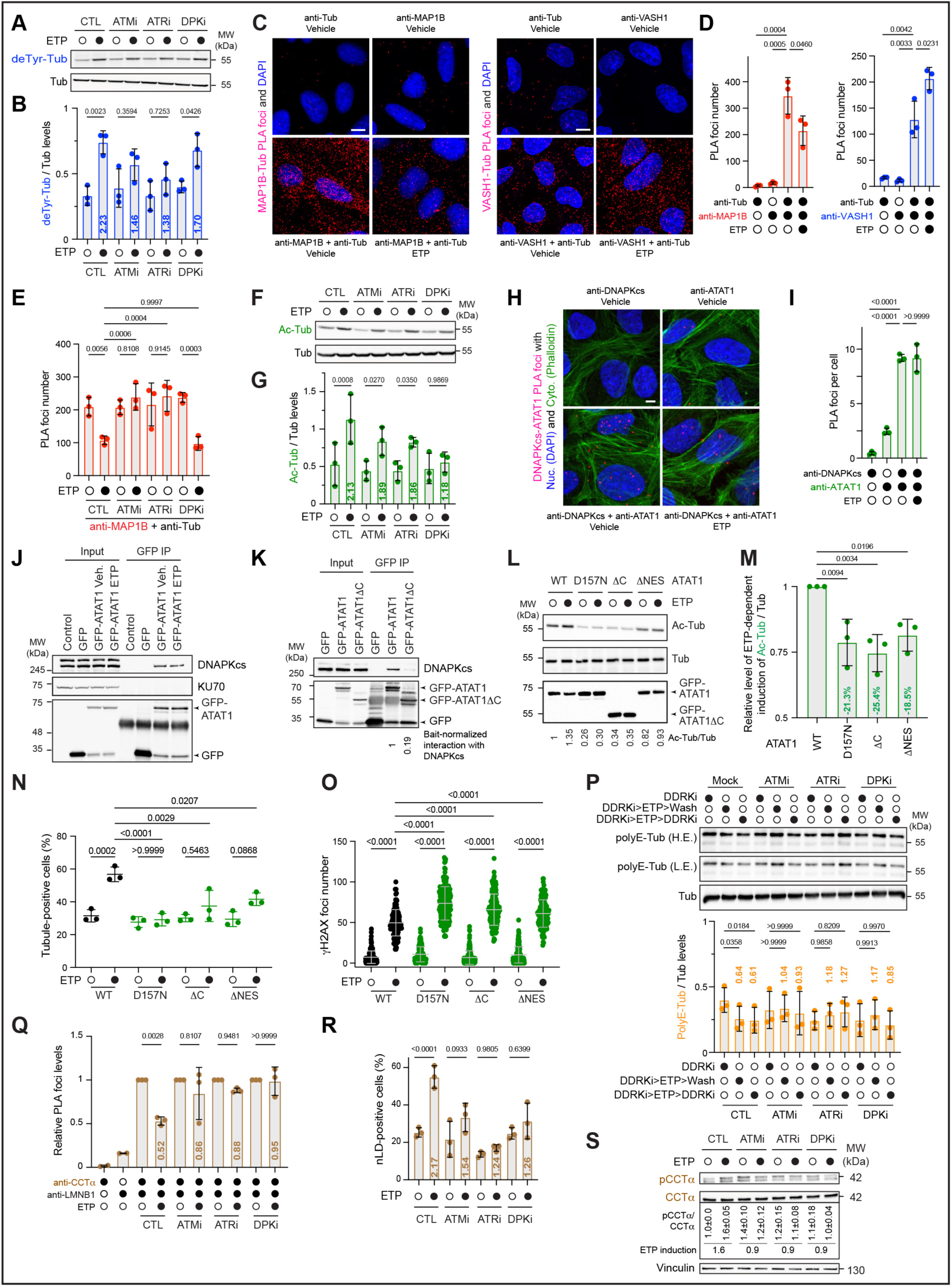
Crosstalk between architectural elements and DNA damage response kinases. (A, B) Immunoblot (A) and quantification (B) of ETP-induced deTyr-Tub following DDR kinase inhibition. (C, D) Representative PLA images (C) and quantification (D) of MAP1B–α-tubulin and VASH1–α-tubulin interactions ± ETP. (E) MAP1B–α-tubulin PLA foci following DDR kinase inhibition. (F, G) Immunoblot (F) and quantification (G) of ETP-induced Ac-Tub following DDR kinase inhibition. (H, I) Representative PLA images (H) and quantification (I) of DNA-PKcs–ATAT1 interactions. (J, K) GFP co-immunoprecipitation of DNA-PKcs with wild-type GFP-ATAT1 (J) or wild-type and C-terminally truncated (ΔC) GFP-ATAT1 (K). (L, M) Immunoblot (L) and quantification (M) of Ac-Tub induction following expression of GFP-ATAT1 constructs ± ETP. (N, O) Percentage of tubule-positive cells (N) and γH2AX foci number (O) in GFP-ATAT1-expressing cells ± ETP. (P) Immunoblot and quantification of polyE-Tub following DDR kinase inhibition, ETP treatment, and recovery. (Q–S) CCTα–LMNB1 PLA foci (Q), nLD frequency (R), and pCCTα levels (S) following DDR kinase inhibition ± ETP. (A-S) Experiments were performed in U2OS cells (A–I and L–S) or HEK293T cells (J and K); Cells were pretreated with ATMi (5 µM, 1 h), ATRi (1 µM, 1 h), or DNA-PKi (20 µM, 2 h), followed by 100 µM ETP unless otherwise indicated; In P, inhibitors were maintained or removed during a 2 h recovery; GFP-ATAT1 constructs (L–O) included wild-type, catalytically inactive (D157N), ΔC, and nuclear export signal deletion (ΔNES) expressed in cells with reduced endogenous ATAT1 (Figure S3D); Tubulin PTMs were normalized to α-tubulin; Data are mean ± SD, n = 3 (A–I, M–S), or 1 (J,K) biological replicates. One-way ANOVA with Dunnett’s (M) or two-way ANOVA with Tukey’s (B, D, E, G, I, N–R) multiple comparisons test; Scale bars, 5 µm (C, H).

Ac-Tub exhibited a distinct DDR kinase dependency, with the strongest dependence on DNA-PKcs and intermediate dependence on ATM and ATR (**Figures 3F and 3G**). Proximity and co-immunoprecipitation assays revealed that ATAT1 associated with DNA-PKcs in both the nucleus and cytoplasm irrespective of etoposide treatment (**Figures 3H–3J and S3B**), while analysis of the ATAT1 interactome using mass spectrometry datasets^45^ independently supported a constitutive ATAT1–DNA-PKcs interaction (**Figure S3C**). Structurally, ATAT1 comprises a structured N-terminal catalytic domain and a disordered C-terminal region, the latter of which is dispensable for intrinsic catalytic activity^46,47^. However, phosphorylation within the C-terminal region disrupts ATAT1’s intracellular localization and consequently its activity^47–49^. Deletion of this C-terminal region of ATAT1 markedly reduced association with DNA-PKcs (**Figure 3K**).

Therefore, we asked how ATAT1 catalytic activity, its C-terminus, and nucleocytoplasmic distribution contribute to damage-induced remodeling. We knocked out endogenous ATAT1 using promoter-targeting sgRNA/Cas9 and expressed wild-type ATAT1, the catalytically inactive D157N mutant, a C-terminal truncation mutant, or a mutant lacking the nuclear export sequence (**Figures S3D–S3F**)^50^. Unlike wild-type ATAT1, all three mutants failed to support etoposide-induced Ac-Tub accumulation and dsbNET formation and resulted in increased γH2AX levels (**Figures 3L–3O**). Thus, damage-induced Ac-Tub and nucleomorphosis require ATAT1’s catalytic activity as well as regulatory features controlling nucleocytoplasmic distribution and interaction with DNA-PKcs.

We next asked whether DDR kinase activity also establishes the polyE-Tub and membrane-remodeling states identified herein. During recovery from a 1-h etoposide pulse, the relative decrease in polyE-Tub depended on ATM, ATR, and DNA-PKcs activity (**Figure 3P**). Similarly, damage-induced decreases in CCTα–LMNB1 proximity at the nuclear periphery, accumulation of nLDs, and phosphorylation of CCTα depended on DDR kinase activity (**Figures 3Q–3S**). Thus, DDR kinases regulate not only individual tubulin modifications, but also architectural states distributed across both cytoskeletal and nuclear membrane compartments.

Collectively, our findings identify DDR kinases as upstream regulators of an emerging architectural code. Distinct kinase dependencies establish different tubulin states, while DDR kinase activity also controls association of CCTα with the nuclear lamina as well as nLD abundance. DNA damage signaling therefore generates the architectural code through coordinated but mechanistically distinct regulation of its cytoskeletal and membrane elements.

### Architectural code elements are functionally integrated

To determine whether architectural code elements function independently or as components of an integrated system, we examined relationships within and between the cytoskeletal and membrane remodeling processes. Although VASH1 and VASH2 both contributed to deTyr-Tub accumulation, VASH2 uniquely promoted baseline and damage-induced Ac-Tub accumulation (**Figure 4A**), consistent with partial functional divergence between VASH1 and VASH2^51,52^. Conversely, disruption of ATAT1/Ac-Tub increased deTyr-Tub in the absence or presence of DNA damage (**Figure 4A**). Thus, loss of ATAT1/Ac-Tub increases deTyr-Tub, whereas loss of VASH2/deTyr-Tub decreases Ac-Tub, revealing bidirectional coupling between tubulin code elements. Together, our observations reveal cross-regulation among tubulin code elements and show that their responses to DNA damage are not independent.

**Figure 4.**
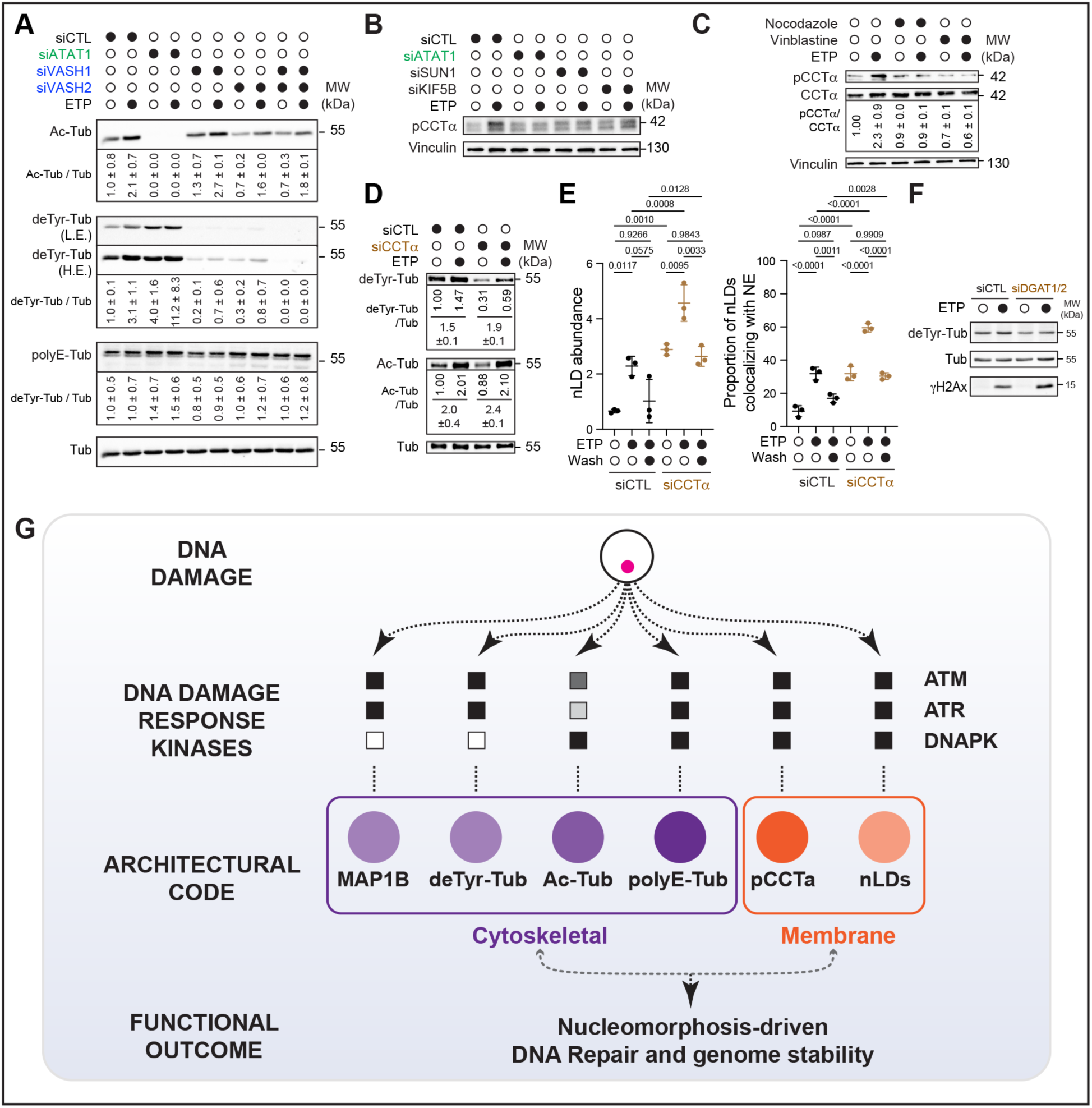
Functional integration of architectural code elements. (A) Immunoblots of tubulin PTMs following ATAT1, VASH1, or VASH2 depletion ± ETP. (B,C) Immunoblot analysis and quantification of pCCTα following ATAT1, SUN1, or KIF5B depletion ± ETP (B), or intact microtubule disruption with 50 µM Nocodazole or 100 nM Vinblastine for 1h ± ETP (C). (D) Immunoblot analysis and quantification of deTyr-Tub and Ac-Tub following CCTα depletion. (E) nLD abundance and colocalization with the nuclear envelope following CCTα depletion, ETP treatment, and recovery (Wash). (F) Immunoblot analysis of deTyr-Tub following combined DGAT1/DGAT2 depletion ± ETP. (G) Proposed architectural code regulating DNA repair. (A-F) Data are mean ± SEM (A, C, D) or mean ± SD (E); n = 3 (E) or 2 (A, C, D) or n=1 (B, F) biological replicates; Two-way ANOVA with Tukey’s multiple comparisons test (E).

Integration of architectural code elements also extended between the cytoskeletal and membrane layers. Damage-induced, but not baseline, CCTα phosphorylation showed dependency on ATAT1, SUN1, KIF5B, and intact microtubules (**Figures 4B and 4C**), linking CCTα regulation to the cytoskeletal machinery that drives damage-induced dsbNET formation^17^. Reciprocally, CCTα depletion decreased deTyr-Tub without altering Ac-Tub (**Figure 4D**) and increased nLD abundance and colocalization with dsbNETs both in the absence and presence of etoposide (**Figure 4E**). Disruption of nLD biogenesis through DGAT1/2 depletion similarly decreased deTyr-Tub (**Figure 4F**). Thus, perturbations originating within either the cytoskeletal or membrane layer altered code elements in the other.

Collectively, our findings indicate that an integrated architectural regulatory system spanning cytoskeletal and membrane layers, rather than a collection of independent processes, drives nucleomorphosis-dependent DNA repair. Architectural elements exert distinct positive and negative effects on damage-induced nuclear remodeling, cross-regulate within each layer, and communicate reciprocally between the layers, allowing regulatory information to propagate across distinct cellular compartments.

## DISCUSSION

Our findings support a model in which DNA damage is interpreted through a distributed architectural code directing nucleomorphosis-driven DNA repair (**Figure 4G**). Unlike classical signaling pathways, in which information is transmitted primarily through sequential molecular interactions, the architectural code described here distributes information across cellular structures themselves. DNA damage triggers coordinated cytoskeletal and membrane reorganizations, guiding nucleomorphosis and DSB repair outcomes.

Several observations support this interpretation. First, DNA damage triggers coordinated, temporally organized transitions in multiple architectural features rather than isolated or independent remodeling events. Second, DDR kinases establish distinct code states through dedicated regulatory factors. Third, code elements are functionally integrated: tubulin modifications cross-regulate while cytoskeletal and membrane code layers communicate reciprocally. Fourth, individual code elements are not functionally equivalent. Specifically, Ac-Tub/ATAT1 and deTyr-Tub/VASH1/2 support damage-induced remodeling and promote HR and NHEJ, whereas the membrane-associated and phosphoregulated CCTα layer licenses nucleomorphosis and alters DDR kinase activation and the relative reliance on different repair pathways. Thus, the biological meaning of the architectural code emerges not from any individual structural feature, but from the combination, temporal organization, and functional integration of multiple architectural states.

Molecular codes are traditionally viewed as information encoded on discrete biochemical substrates, such as histones or the RNA polymerase II C-terminal domain^7–10^. In contrast, the architectural code described here distributes information across components of cytoskeletal and membrane layers. Therefore, we propose that the components and regulators of cellular architecture can collectively function as an information-bearing substrate, integrating biochemical signaling with large-scale structural responses critical to genome stability. Architectural code elements communicate across cellular compartments and exert distinct effects on the DDR. Such distributed organization may represent a broader strategy through which cells convert molecular signals into coordinated cellular behaviors.

Nucleomorphosis provides a higher-order structural output of the architectural code. Our previous work identified dsbNETs as transient nuclear-envelope structures that facilitate DSB repair^17^. Here, we place this response within a broader regulatory system in which the tubulin modifications, cytoskeletal machinery, CCTα regulation, and nuclear lipid remodeling are functionally integrated. Rather than representing an isolated morphological response to DNA damage, nuclear reshaping therefore emerges from coordinated regulation across cytoplasmic and nuclear-envelope structures. In this framework, the architectural code controls when and to what extent nucleomorphosis occurs while coupling nuclear remodeling to distinct DNA repair outcomes.

Our work also raises a number of intriguing questions. Although we identify functionally integrated cytoskeletal and membrane code elements and their regulators, the processes through which code units are established, propagated, functionally interpreted, or even possibly inherited will require future studies. Such efforts may eventually identify a much larger network of factors that write, read, and execute architectural code states, expanding our understanding of the molecular mechanisms underlying communication between the DDR kinases, cytoskeleton, and nuclear envelope and lipids. Additionally, DNA damage-induced CCTα inactivation, nLD remodeling, and cPLA2 perinuclear recruitment suggest that nuclear membrane lipid composition transiently changes to accommodate mechanical forces on the nuclear envelope. Future work could explore such potential changes under diverse DNA-damaging conditions. It is also intriguing to speculate that post-exposure decreases in polyE-Tub persist and influence cellular response upon re-exposure to DNA damage. Future studies may also provide additional insight as to how the unique activation of individual code elements influences specific DNA repair steps and pathways differently across the genome. Finally, it will also be exciting to explore whether similar architectural coding principles operate during genome maintenance processes beyond DSB repair, or during cellular responses to mechanical or environmental signaling cues that reflect mechanical stress triggered by developmental or pathological changes occurring at the tissue or organ level.

Collectively, our findings identify a distributed architectural code that connects DNA damage signaling to nucleomorphosis and DNA repair. More broadly, we propose that the broad cell architecture constitutes an information-bearing substrate that is controlled and interpreted by dedicated machinery to transform genome-damaging inputs into distinct genome stability outcomes.

## ACKNOWLEDGEMENTS

We thank Drs. Jeremy Stark and Linda Penn for materials. We thank the groups of Drs. Marco Foiani, Taekjip Ha, Razq Hakem, and Karim Mekhail for discussions. We thank Dr. Paul Paroutis at the SickKids Children’s Hospital Imaging Facility for technical advice. We thank Dr. Jonathan St-Germain at the Princess Margaret Cancer Research Centre for assistance with Mass Spectrometry analysis. A. Hundal is supported by scholarships from the Ontario Graduate Scholarship Program, Peterborough K.M. Hunter Graduate Award, and Canada Graduate Scholarship. D.U. is supported by scholarships from the Ontario Graduate Scholarship Program. R.H. and P.G.M. are supported by Canadian Institutes of Health Research (CIHR) grants (R.H. 183597, 186072; P.G.M. 173542). This research was primarily supported by CIHR grants (180469, 190143) to K.M., who is supported by the Royal Society of Canada.

## AUTHOR CONTRIBUTIONS

A. Hundal, D. Urman, R. Hakem and K. Mekhail conceived the project and designed the experiments. A. Hundal, D. Urman and K. Mekhail wrote the manuscript with input from all authors. A. Hundal and D. Urman designed, conducted, and analyzed imaging, immunoblot, PLA, cloning, and cell cycle assays with help from R. Krishnan for FACS analysis. A. Hundal designed, conducted, and analyzed CoIP and comet assays. A. Hundal designed and conducted DSB repair reporter assays with help from R. Krishnan for FACS analysis. A. Hundal designed and conducted 3D nuclear reconstructions with help from C. A. Jordan and A. Algouneh for imaging and analysis. A. Hundal was assisted by A. Algouneh for imaging and immunoblotting, J. Hao for the irradiation of cells, and A. Hakem for general experimentation. C. A. Jordan and T. Chkuaseli designed and conducted CHEF runs. A. Hundal and D. Urman conducted and analyzed very-fast CRISPR experiments with help from S. Rudnizky and A. T. Rybczynski in designing experiments. P. G. Maass and T. Ha contributed resources and expertise. T. Ha supervised S. Rudnizky and A. T. Rybczynski, while R. Hakem supervised A. Hundal, R. Krishnan, and T. Chkuaseli, and K. Mekhail supervised A. Hundal, D. Urman, A. Algouneh, C. A. Jordan, and T. Chkuaseli. The overall direction and supervision of the project was conducted by K. Mekhail.

## RESOURCE AVAILABILITY

### Lead contact

Further information and requests for resources and reagents should be directed to the lead contact, Karim Mekhail

### Materials, data and code availability

• Materials generated in this study are available upon reasonable request.

• This paper does not report original computer code

## DECLARATION OF INTERESTS

A. Hundal, R. Hakem, and K. Mekhail are listed as inventors on patent applications related to dsbNET-targeting strategies in cancer and aging. The other authors declare no competing interests.

## DECLARATION OF GENERATIVE AI AND AI-ASSISTED TECHNOLOGIES

During the preparation of this work, the authors used ChatGPT and Grammarly for suggestions on how to improve the main text’s readability and grammar. The authors have carefully reviewed and edited any suggested content before adoption and are responsible for the content of the article.

## METHODS

### Cell lines, reagents G treatments

U2OS (ATCC HTB-96), HEK293T (ATCC CRL-3216), DR-GFP U2OS and EJ5 U2OS (gift from Jeremy Stark) cells were cultured in DMEM without sodium pyruvate (Wisent Bioproducts) with 10% v/v FBS (Wisent Bioproducts) and 1% v/v penicillin-streptomycin (Wisent Bioproducts). SUM149PT and HCC1937 (gift from Linda Penn) were cultured in RPMI (Wisent Bioproducts) with 10% v/v FBS and 1% penicillin-streptomycin. Cell lines were frequently tested for mycoplasma with MycoAlert® PLUS Mycoplasma Detection Kit (LT07-218, Lonza).

Unless specified otherwise, cells were treated with 100 µM etoposide (E7657, LKT Laboratories) for 1 h. Pre-treatment of cells with DDR kinase inhibitors was as follows: 5 µM ATM inhibitor (AZD1390, Selleck Chemicals) for 1 h, 1 µM ATR inhibitor (VE-822, Selleck Chemicals) for 1 h, 20 µM DNAPK inhibitor (S2893, Selleck Chemicals) for 2 h before etoposide treatment.

Cells were exposed to γ-irradiation using a Gammacell 40 Exactor Self-Contained Low Dose Rate Research Irradiator (Best Theratronics Ltd.) equipped with a ^137^Cs source. Cells were irradiated at the indicated doses and then returned immediately to standard culture conditions for the specified recovery periods before downstream analyses. The irradiation time was adjusted according to the calibrated dose rate of the instrument at the time of the experiment.

To induce oleate-dependent CCTα responses, cells were incubated in 500 µM BSA-oleate complex (6:1 oleate to BSA molar ratio) (Cayman Chemicals # 29557) for 30 min before subsequent treatments. Pre-treatment of cells with microtubule poisons was as follows: 50 µM Nocodazole (487928-10MG, Sigma-Aldrich; 1228, Tocris) for 1 h, 100 nM Vinblastine (V1377-10MG, Sigma-Aldrich) for 1 h. To deplete lipid droplets, cells were treated with 500 nM Triacsin-C for 24 h.

### Knockdowns G transfections

For transient knockdowns, cells were transfected for 48 h with 50 nM of CCTα-targeting siRNA using Lipofectamine™ RNAiMAX Transfection reagent (137778150, Invitrogen™) according to the manufacturer’s protocol. For the remaining knockdowns, cells were transfected for 72 h with 30 nM of siRNA. All siRNAs used are listed in **Supplementary Table 1**. For transient overexpression of gene constructs, cells were transfected with 2-5 µg of DNA for 72 h with Lipofectamine™ 3000 (L3000015, Invitrogen™) according to the manufacturer’s protocol. Lipofectamine™ 2000 (11668030, Invitrogen™) was used for co-transfection of CCTα-3’UTR-targeting siRNA alongside various CCTα plasmids according to the manufacturer’s protocol. Transfection efficiency was assessed via microscopy or immunoblotting for cells transfected with GFP-expressing constructs. All plasmids used are listed in **Supplementary Table 2**. Knockdowns were validated using various methods, including RT-qPCR, immunoblotting and immunofluorescence as shown.

### vfCRISPR

For immunofluorescence and immunoblotting assays, U2OS cells were seeded in 6-well plates containing coverslips. The next day, cells were transfected with 6.25 µg of recombinant Cas9 and 7.2 µg of duplexed tracrRNA and AluGG cgRNA (Taekjip Ha’s group^40,41^) per well using Lipofectamine™ CRISPRMAX™ (CMAX00015, Invitrogen™) according to the manufacturer’s protocol. To induce the breaks, cells were exposed to UV light for 1 min using the uvBeast™ V3 365nm MINI light UV flashlight. 1 h post-light exposure, coverslips were removed from the 6-well and placed into 24-well dishes for fixation and immunofluorescence analysis, while remaining cells were harvested for immunoblotting.

### Cloning G bacterial transformation

pEF5B-FRT-GFP-ATAT1 ΔC, ΔNES and GFP vector control mutant plasmid constructs were made using the Q5® Site-Directed Mutagenesis Kit (E0554S, New England Biolabs®) according to the manufacturer’s protocol. NEBasechanger® was used to design mutagenesis primers (see **Supplementary Table 3** for the list of primers used).

pcDNA3.1+–PCYT1A/CCTα plasmid was designed and then synthesized (Invitrogen). EGFP was cloned into the plasmid using AgeI and XhoI restriction sites. The CCTα quintuple phosphomimetic mutant (S315D, S319D, S321D, S322D, and S323D) plasmid was generated via two sequential PCRs using mutagenic primers (see **Supplementary Table 3**) and by following the manufacturer’s protocol for site-directed mutagenesis (New England Biolabs®). The quintuple phosphonull mutant (S315A, S319A, S321A, S322A, and S323A) fragment was custom-ordered from Invitrogen™ and cloned into pcDNA3.1+–PCYT1A/CCTα plasmid via in-gel ligation. Briefly, 1.5 µg of pcDNA3.1+–PCYT1A/CCTα plasmid and 3 µg of the phosphonull mutant fragment were digested with AgeI-HF (R3552S, New England Biolabs) and EcoRI (R3101S, New England Biolabs) at 37°C overnight. Digested DNA products were run on a 0.7% low-melting point agarose gel at 100 V for 1 h at 4°C. Bands were imaged with a UV transilluminator and excised per the expected size. The excised bands were melted at 70°C for 10 min and used at a 1:3 vector-to-insert ratio. The ligation reaction was performed using T4 DNA Ligase (M0202S, New England Biolabs) at 16°C overnight. The reaction was heat-inactivated at 65°C, and the products were transformed into DH5*a* competent *E. coli*.

DH5*a* competent *E. coli* cells were transformed with 1 µg plasmid for 30 min on ice. Bacteria were heat-shocked at 42°C for 45 seconds and then immediately transferred to ice for 2 min. Then, cells were revived in Luria-Bertani (LB) broth for 1 h, rotating at 37°C. Cells were centrifuged at 14,000 × *g* for 2 min, resuspended in 100 µL of LB, and then plated onto LB plates with the appropriate selection antibiotic at 37°C overnight. Colonies were picked the next day and grown overnight in LB + antibiotic (125 µg/mL Ampicillin) medium. Plasmids were isolated from cultures according to the manufacturer’s protocol (GeneAid).

### Knockout G lentiviral preparation

A gRNA sequence targeting the promoter region of ATAT1 was designed using CHOPCHOP-v3^53^ (see **Supplementary Table 1**). The gRNA was cloned into pLentiCRISPR v2 (Addgene) using a described protocol^54,55^. Stbl2™ cells (Invitrogen™) were transformed with the plasmid and grown at 30°C overnight on LB plates with Ampicillin selection. Positive colonies were confirmed by sequencing. HEK293T cells were transfected with 4 µg of plentiCRISPR-v2 plasmid (52961, Addgene) containing the ATAT1 gRNA, 4 µg of psPAX (12260, Addgene), and 2 µg pMD2.G (12259, Addgene) using GeneJet™ In Vitro Transfection Reagent (SL100499 SignaGen Laboratories®) according to the manufacturer’s protocol. The virus was harvested 72 h post-transfection using a 0.2 µm sterile filter and mixed with Polybrene to a final concentration of 8 µg mL^-1^. U2OS cells were transduced with the virus for 24 h, recovered in DMEM with 10% v/v FBS and 1% v/v penicillin-streptomycin for 24 h, and selected using 2 µg mL^-1^ puromycin for 7 d. Knockout was confirmed via immunoblotting.

### RNA extraction G RT-qPCR

Cells grown to 70-80% confluence were washed with RNase-free PBS before RNA extraction using the RNeasy Kit (74104, Qiagen) according to the manufacturer’s instructions. Isolated RNA was quantified, and 1 µg of RNA was treated with DNase I Amp Grade (1 U µl^−1^; 18068015, Thermo Fisher Scientific) for 15 min at 23°C. The reaction was quenched with 25 mM EDTA and incubated at 65°C for 10 min. cDNA synthesis was performed by reverse transcription, using 10 µL of RNA, 10 mM dNTPs, 1 µL random hexamer primers, 100 mM DTT, 5x First-Strand buffer, 200 U/µL M-MLV reverse transcriptase (28025013, Invitrogen), and 40 U/µL RNaseOUT (10777019, Invitrogen) according to the manufacturer’s instructions. PCR conditions were 10 min at 25°C, 1 h at 37°C, and 15 min at 70°C. The resulting cDNA was diluted 1:10, and 4 µL of cDNA was used per sample for qPCR. Wells were topped up to 10 µL with final reaction consisting of 1x SensiFAST SYBR No-ROX Mix (BIO-98050, Bioline) and 400 nM each of primers. The qPCR reaction was performed at 95°C for 2 min, 60°C for 30 sec, followed by 39 cycles of 95°C for 5s and 60°C for 30s. Relative mRNA levels were quantified using the formula: 2^!ΔΔ*Ct*^, *w*ℎ*ere* ΔΔ*Ct* = Δ*Ct*(*siTarget*) −Δ*Ct*(*siControl*), *w*ℎ*ere* Δ*Ct* = *Ct*(*transcript of interest*) − *Ct*(ℎ*ousekeeping transcript*)

### Immunofluorescence

Cells seeded on glass coverslips were fixed with 4% v/v paraformaldehyde (PFA) in PBS or 100% ice-cold methanol for 10 min at room temperature (RT) or 4°C, respectively. When probing for pS4/pS6-RPA2, cells were briefly incubated in cytoskeleton (CSK) buffer (20 mM HEPES pH 7.0, 100 mM NaCl, 3 mM Sucrose, 3 mM MgCl_2_, 0.5% Triton X-100) for 5 min on ice or 4°C prior to fixation with 4% PFA. Coverslips fixed with 4% v/v PFA were permeabilized with 0.5% v/v Triton X-100 in PBS at 23°C for 10 min. Coverslips were then blocked with 5% w/v bovine serum albumin (BSA) in PBS at 23°C for 1 h. Coverslips were transferred to a humid, light-proof chamber. Primary antibodies diluted in 1% w/v BSA were added to the coverslips (all antibodies used with their dilutions are listed in **Supplementary Table 4**). Coverslips were incubated in primary antibodies at 4°C overnight. Next, coverslips were washed 3 × 5 min with PBS. Secondary antibodies diluted in 1% BSA w/v were added to the coverslips and incubated at 23°C for 1 h. Secondary antibodies were washed with PBS 5 × 5 min at RT. Coverslips were incubated in DAPI (D1306, Thermo Fisher Scientific) at 23°C for 5 min. For lipid droplet visualization, coverslips were then stained with BODIPY493/503 (4,4-difluoro-1,3,5,7,8-pentamethyl-4-bora-3a,4a–diaza-s-indacene; D3922, Thermo Fisher Scientific) diluted in PBS at 23°C for 10 min. Coverslips were briefly washed with PBS and then mounted on imaging slides using Mowiol (81381, Sigma-Aldrich) mounting medium. Slides with mounted coverslips were left at 23°C overnight for the Mowiol to set prior to imaging the next day or storage at 4°C.

### Fixed-cell confocal microscopy

The Nikon Eclipse Ti2 C2+ coupled to NIS-Elements AR software (Nikon, version 4.50.00) or the Leica Stellaris 5 WLL confocal microscope coupled to Leica LASX software (version 4.8.2.29567) were used to acquire images. 3D images were acquired by imaging z-stacks with a maximum step size of 0.5 µm using a 60x oil immersion objective. Super-resolution imaging was performed using the Zeiss LSM880 Airyscan confocal microscope with the Zen Black software (version 2.3) and SR mode with a maximum step size of 0.2 µm using a 63x oil immersion objective as described^17^.

### Image analysis

#### Nuclear reconstruction

Nuclear reconstructions were performed using BitPlane Imaris version 10.1.1. The Imaris ‘Surfaces’ function with automatic segmentation was used to create a peripheral nuclear boundary surface using the DAPI channel. To segment the internal Lamin-B1 tubule structures, the ‘Surfaces’ function with machine learning segmentation was used. Internal lamin-B1 structures were labeled using the Lamin-B1 channel to train the classifier. The ‘Cells’ function was used to integrate the boundary and internal segmentations as previously described^17,56^. The surface areas were calculated using the ‘detailed statistics’ function, and the tubule score was calculated by dividing the total nuclear surface area (boundary+internal) by the boundary surface area.

#### dsbNETs analysis

Single-cell analysis was performed using either BitPlane Imaris (version 10.1.1) or ImageJ/FIJI (version 2.16.0). Lamin-B1-stained nuclei were analyzed for dsbNETs as previously described^17^. For 3D visualization of the nuclear lamina in ImageJ, the “orthogonal views” function was used to visualize tubules in the xz and yz planes. Nuclei were considered dsbNETs positive if two or more tubules traversed at least half the distance of the nuclear diameter^17^.

#### Foci and lipid droplet analyses

ImageJ was used for single-cell analysis of DNA damage foci and nuclear lipid droplets. Maximum projection images were generated and analyzed using the ‘Difference of Gaussians’ and threshold and “Analyze Particles” functions.

### Proximity ligation assays

Duolink® PLA was performed according to the manufacturer’s protocol (Sigma-Aldrich). Briefly, coverslips were fixed and permeabilized as described above. Then, coverslips were blocked with Duolink Blocking Buffer at 37°C for 1 h. Coverslips were incubated at 4°C overnight with primary antibodies diluted 1:500 v/v in the antibody diluent. Coverslips were washed with the provided Wash Buffer A, followed by incubation with PLA probes (Anti-Rabbit PLUS and Anti-Mouse MINUS) at 37°C for 1 h. Duolink *in situ* Detection Reagents Red (DUO92101, Sigma-Aldrich) were used to perform the ligation and amplification reactions, where coverslips were incubated at 37°C for 30 min and 100 min, respectively. Finally, coverslips were washed, incubated in DAPI solution at 23°C for 5 min, and then mounted onto imaging slides with Mowiol mounting reagent. Slides were imaged within the next 1-2 d for optimal PLA signal capture. Slides were kept at -20°C for longer-term storage.

### SDS-PAGE G Immunoblotting

Cells were harvested with cell scrapers and lysed with RIPA buffer (10 mM Tris-HCl pH 8.0, 1 mM EDTA, 0.5 mM EGTA, 1% Triton X-100 (v/v), 0.1% sodium deoxycholate (w/v), 0.1% SDS (w/v), 140 mM NaCl, cOmplete™ protease inhibitor (Roche)) on ice for 1 h. Lysates were centrifuged at 8,000 × *g* at 4°C for 10 min to remove debris. The Bradford assay (Bio-Rad) was used to determine protein concentration. Samples were diluted to equal protein concentration and boiled in 3X Laemmli buffer (187.5 mM Tris-HCl pH 6.8, 6% SDS (w/v), 30% glycerol (v/v), 150 mM dithiothreitol, 0.03% bromophenol blue (w/v)) at 95°C for 5 min. Samples were run on 6-12.5% SDS-polyacrylamide gels or 4-20% Novex™ Tris-Glycine Mini Protein Gel Kits (XP04200BOX, Invitrogen™) depending on the probed protein size. Proteins were transferred onto nitrocellulose membranes and then blocked in 5% skim milk/TBST or 5% BSA/TBST (w/v) at 23°C for 1 h. Primary antibodies diluted in 5% BSA/TBST (w/v) were added to the blots and incubated at 4°C overnight. The following day, membranes were washed using 3 × 5 min TBST at 23°C. HRP-conjugated secondary antibodies diluted in blocking buffer were added to the blots and incubated at 23°C for 1 h. Blots were washed using 3-5 × 10 min TBST at 23°C. Blots were incubated with ECL substrate (Clarity Western ECL Substrate, Bio-Rad) for 5 min and developed using the ChemiDoc imaging system (Bio-Rad). Blots were quantified using Image Lab (version 6.0.1).

### Co-immunoprecipitation

HEK293T cells were transfected with 2µg GFP vector, 5 µg wild-type GFP-ATAT1, and 10 µg ΔC mutant GFP-ATAT1, and harvested 48 h post-transfection. Cells were lysed on ice for 1 h with RIPA buffer and centrifuged at 8,000 × *g* for at 4°C for 10 min to remove debris. Lysates were pre-cleared in 10 µL Dynabeads™ Protein G magnetic beads (10003D, Invitrogen™) at 4°C for 1 h with rotation. Meanwhile, 3 µg of GFP antibody was incubated with 30 µL beads in 500 µL IP buffer (10 mM Tris-HCl pH 7.4, 1 mM EDTA, 1 mM EGTA, 150 mM NaCl, 0.5% NP-40 (v/v), 0.2 mM sodium orthovanadate, cOmplete™ protease inhibitor) at 4°C for 1-2 h with rotation. Antibody-conjugated beads were washed with IP buffer using a magnetic rack at 23°C. A portion of the pre-cleared lysates was retained as input, and the rest were added to the conjugated beads and incubated at 4°C for 2 h with rotation. Beads were washed with IP buffer at 4°C for 3 × 5 min with rotation and then quickly washed twice at 23°C. Proteins captured by the conjugated beads were eluted by adding 50 µL of 1X Laemmli buffer and incubated at 95°C for 5 min. Beads were removed, and IPs were transferred to fresh tubes and stored at -80°C.

### Cell cycle analysis

Cells were grown to 70-80% confluence and incubated in 10 µM EdU (5-ethynyl-2’-deoxyuridine) in DMEM at 37°C for 1 h. Cells were trypsinized, washed with PBS, and EdU detection was achieved with the Click-iT™ plus EdU cell Pacific Blue™ Flow Cytometry Assay kit (C10636, Invitrogen™) according to the manufacturer’s protocol. The samples were treated with RNase A and stained with propidium iodide to label total DNA content. The measurements were acquired using a BD LSR Fortessa flow cytometer, and data were analyzed using FlowJo software (version 10).

For immunofluorescence-based analysis, cells were incubated in 10 µM EdU in DMEM at 37°C for 1 h. Cells were fixed with 4% PFA at 23°C for 10 min and permeabilized using 0.5% Triton X-100 (v/v) at 23°C for 10 min. The Click-iT reaction was performed prior to blocking using the Click-iT™ Plus EdU Cell Proliferation Kit for Imaging, Alexa Fluor™ 488 dye (C10637, Invitrogen™) according to the manufacturer’s protocol. After the Click-iT™ reaction, coverslips were washed in PBS three times before proceeding with blocking and the subsequent immunofluorescence protocol.

### Neutral comet assay

Experiments were performed using the Comet Assay Kit (ab238544, Abcam) according to the manufacturer’s instructions. Cells were treated with DMSO or 100 µM etoposide and harvested immediately following a 1 h treatment. For the wash condition, cells were treated with 100 µM etoposide for 1 h, washed with PBS, and recovered in DMEM for 45 min prior to harvesting. Subsequent steps were performed as previously described^17^. Data were analyzed using CometAnalyser (version 1.2.0)^57^.

### Nuclear-cytoplasmic fractionation

To achieve comparable expression levels, U2OS cells were transfected with either 2.5 µg of wild-type GFP-ATAT1, 2.5 µg GFP-ATAT1-D157N, 5 µg GFP-ATAT1-ΔC, or 5 µg GFP-ATAT1-ΔNES. Subcellular fractionation was performed using the NE-PER™ Nuclear and Cytoplasmic Reagents (78833, Thermo Scientific™) according to the manufacturer’s protocol. Briefly, cell pellets were resuspended in ice-cold CERI buffer, incubated on ice for 10 min before the addition of ice-cold CERII buffer, and then centrifuged at 16,000 × *g* for 5 min. The supernatants were retained as the cytoplasmic fractions, and the remaining pellets containing nuclei were resuspended in ice-cold NER buffer and incubated on ice for 40 min. The samples were then centrifuged at 16,000 × *g* for 10 min, and the resulting supernatant was retained as the nuclear fraction. The CER1:CERII:NER ratio was 200:11:100 µL.

### HR G NHEJ reporters

DR-GFP (HR) and EJ5 (NHEJ) cells were reverse-transfected with 30 nM siRNA using RNAiMAX (13778150, Invitrogen™). The following day, cells were transfected with 3 µg I-SceI plasmid (26477, Addgene) or empty vector control (pCAGGS-BSKX, gift from Jeremy Stark). 48 h post I-SceI transfection, cells were trypsinized, washed in PBS, and resuspended in 250 µL sorting buffer (1 mM EDTA, 25mM HEPES pH 7.0, 1% Fetal Bovine Serum in PBS). Cells were run on the BD LSR Fortes cell analyzer and analyzed using FlowJo software (version 10).

### CHEF analysis

The CHEF was performed with modifications from an established protocol^58^. Briefly, cells were treated with etoposide for the indicated time, and 5×10^5^ cells were harvested and resuspended in PBS. 45 <u>µL of</u> cells resuspended in PBS was mixed with 45 µL of prewarmed 1.2% (w/v) low-melting-point agarose in PBS. The cell and agarose mixture was solidified as agarose plugs in plug moulds (Cat. Nb. 1703713, Bio-Rad). Plugs were incubated in NDSK buffer (0.5 M (w/v) EDTA, 10 mg ml^-1^ N-lauryl sarcosine (w/v), 1mg ml^-1^ proteinase K) at 50°C for 48 h. Plugs were then incubated with 1X cOmplete, EDTA-free protease inhibitor (Roche) in 1X TE at 23°C for 24 h with gentle shaking, followed by an additional incubation in 1X TE at 4°C for 24 h. Subsequently, plugs were incubated with 50 units of NotI-HF in 1X rCutsmart buffer (New England Biolabs®) at 37 °C for 24 h to digest DNA. The plug samples were then run on a 1% contour-clamped homogeneous field electrophoresis (CHEF)-specific agarose gel and resolved in 1X TAE using a CHEF DR II apparatus (Bio-Rad, United States) for 48 h (3 V cm^-1^, 0.2-266 sec). The gel was stained with ethidium bromide for DNA visualization before imaging using the ChemiDoc imaging system (Bio-Rad).

### Statistics G reproducibility

Statistical analysis was performed using GraphPad Prism software using the statistical tests as described. Data were generated using the indicated number of biological replicates. Immunoblots and microscopy images are representative of at least two independent biological replicates unless indicated otherwise.

