## Supplementary Information for "A Distributed Architectural Code Governs DNA Repair"

**Supplementary Information Content:**

Supplementary Figures 1-3
Supplementary Tables 1-4

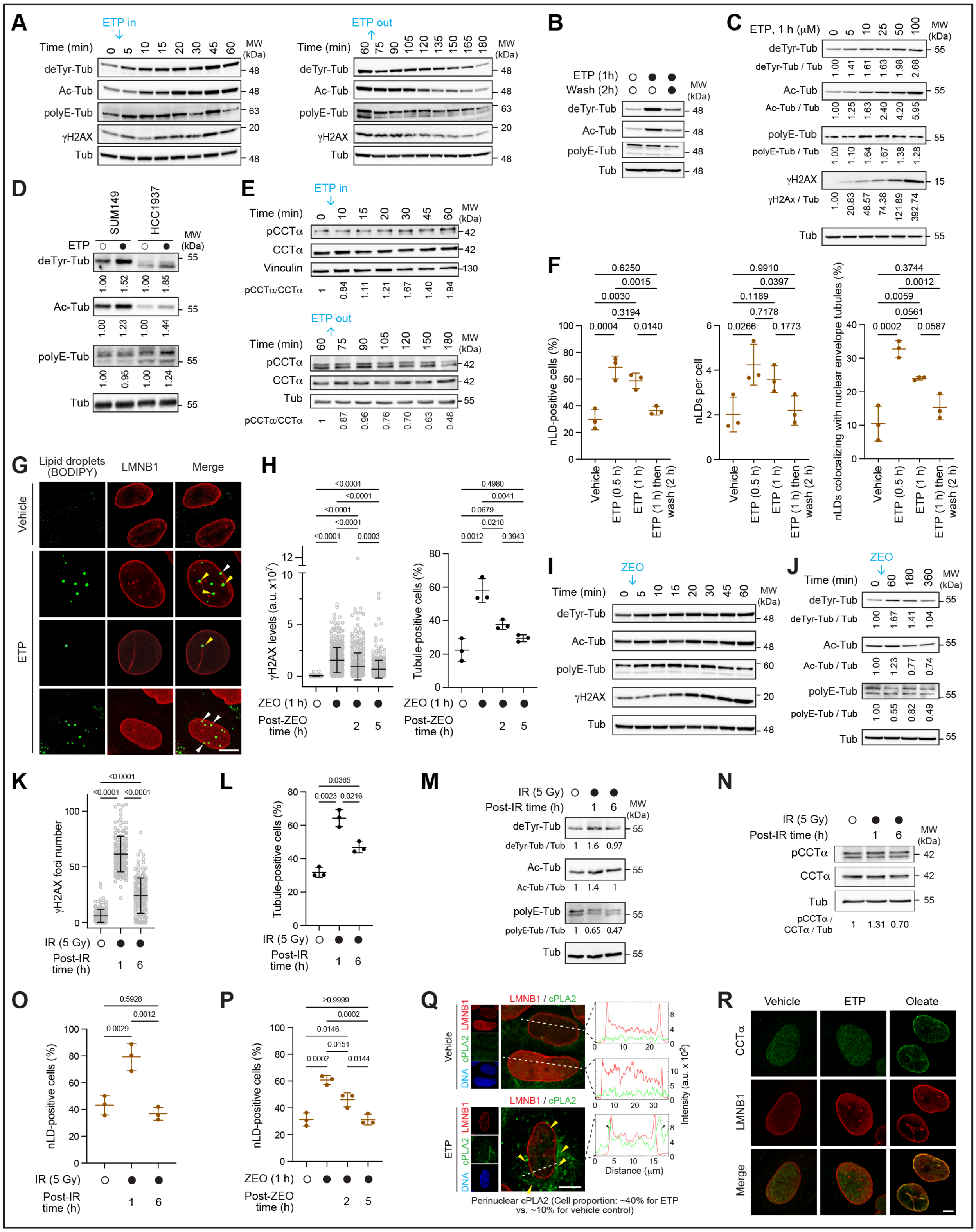

**Supplementary Figure 1. Architectural reconfiguration under various DNA-damaging agents and contexts; related to Figure 1.** (A, B) Representative immunoblots of tubulin PTMs during the treatment and recovery from etoposide (ETP) treatment corresponding to

Figure 1. (C) Immunoblot analysis of tubulin PTMs and γH2Ax following increasing ETP concentrations. (D) Immunoblot of tubulin PTMs in BRCA1-mutant SUM149PT and HCC1937 cells ± ETP. (E) Representative immunoblot of pCCTα induction and recovery corresponding to Figure 1. (F, G) Quantification (F) and representative images (G) of nLD frequency, abundance, and nuclear envelope colocalization. White and yellow arrows indicate nLD contacts with the nuclear periphery and nuclear envelope tubules, respectively. (H–J) γH2AX nuclear intensity, percentage of tubule-positive cells (H), and tubulin PTM levels (I, J) following Zeocin treatment and recovery. (K–N) γH2AX foci number (K), percentage of tubule-positive cells (L), tubulin PTM levels (M), and pCCTα levels (N) following ionizing radiation (IR). (O, P) nLD frequency following IR (O) or Zeocin treatment (P). (Q) Representative images and intensity profiles of perinuclear cPLA2 localization. Yellow arrows indicate perinuclear cPLA2 enrichment; black arrows indicate cPLA2 intensity peaks adjacent to LMNB1 peaks that indicate the nuclear periphery. (R) Representative images of CCTα localization following vehicle, ETP, or oleate treatment. (A-R) Treatments included ETP (100 μM, 1h), Zeocin (400 μg/mL, 1 h), IR (5 Gy), and oleate (500 μM, 30 min), with recovery durations as indicated where applicable; Immunoblot signals were normalized to loading controls and untreated conditions; Data are mean ± SD; n = 3 (A, B, C, F, H, I, K, L, O, P) or 2 (D, E, J, M, N) biological replicates; Two-way ANOVA with Tukey's multiple comparisons test (F, H, K, L, O, and P); Scale bars, 10 μm (G, Q) and 5 μm (R).

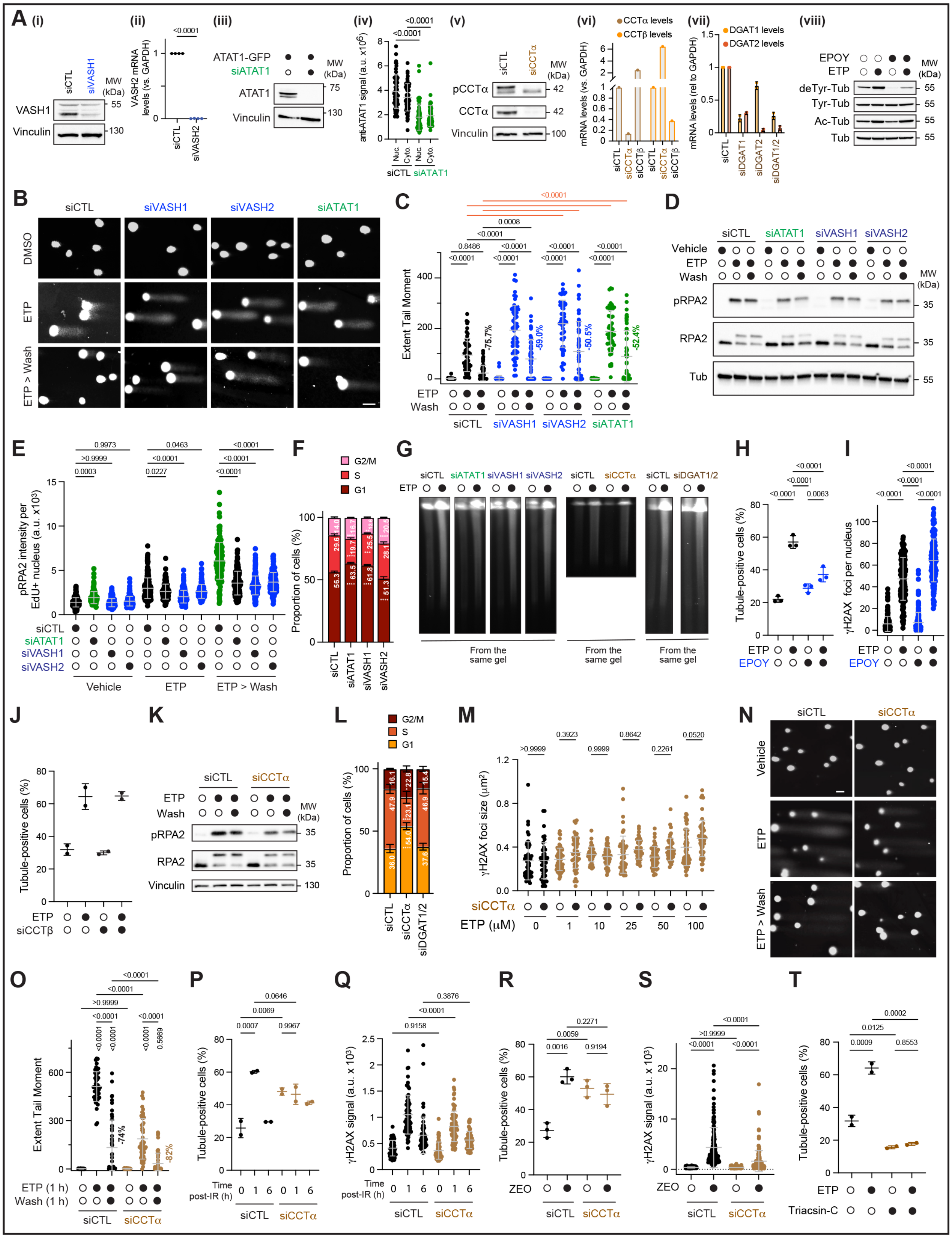

**Supplementary Figure 2. Architectural elements provide distinct contributions to DNA repair; related to Figure 2.** (A) Validation of VASH1, VASH2, ATAT1, CCTα, CCTβ, DGAT1, and DGAT2 depletion by immunoblotting, RT-qPCR, or immunofluorescence, as indicated, and immunoblot analysis of tubulin PTMs following vasohibin inhibition with EPOY. (B, C) Representative neutral comet assay images (B) and extent tail moment quantification (C) following VASH1, VASH2, or ATAT1 depletion, etoposide (ETP) treatment, and recovery. (D, E) Immunoblot analysis of phospho-RPA2 (Ser4/Ser6) and total RPA2 (D), and phospho-RPA2 nuclear intensity (E), following ATAT1, VASH1, or VASH2 depletion, ETP treatment and 4h recovery. (F) Cell-cycle distribution following ATAT1, VASH1, or VASH2 depletion. (G) CHEF gel analysis of DNA fragmentation following ATAT1, VASH1, VASH2, CCTα, or combined DGAT1/DGAT2 depletion ± ETP for 20 min. (H, I) Percentage of tubule-positive cells (H) and γH2AX foci number (I) following vasohibin inhibition with EPOY. (J) Percentage of tubule-positive cells following CCTβ depletion ± ETP. (K) Immunoblot analysis of phospho-RPA2 and total RPA2 following CCTα depletion, ETP treatment, and 4h recovery. (L) Cell-cycle distribution following CCTα depletion or combined DGAT1/DGAT2 depletion. (M) γH2Ax foci size following CCTα depletion and treatment with the indicated ETP concentrations. (N, O) Representative neutral comet assay images (N) and extent tail moment (O) following CCTα depletion, ETP treatment, and 45 min recovery. (P–S) Percentage of tubule-positive cells (P, R) and γH2Ax foci intensity (Q, S) following ionizing radiation (IR) treatment (P, Q) or Zeocin (ZEO) treatment (R, S) in CCTα-depleted cells. (T) Percentage of nuclear envelope tubule-positive cells following ETP treatment with or without Triacsin C treatment. (A-T) Cells were pretreated with EPOY (20 μM, 2 h) or Triacsin C (0.5 μM, 24 h) where indicated; Data are mean ± SD; n = 3 (E, F, H, I, L, M, R, S) or 2 (D, J, K, T) biological replicates; n = 50 comets per condition (C, O); Two-way ANOVA with Tukey's multiple comparisons test (C, E, F, H, I, M, O, P, Q, R, S).

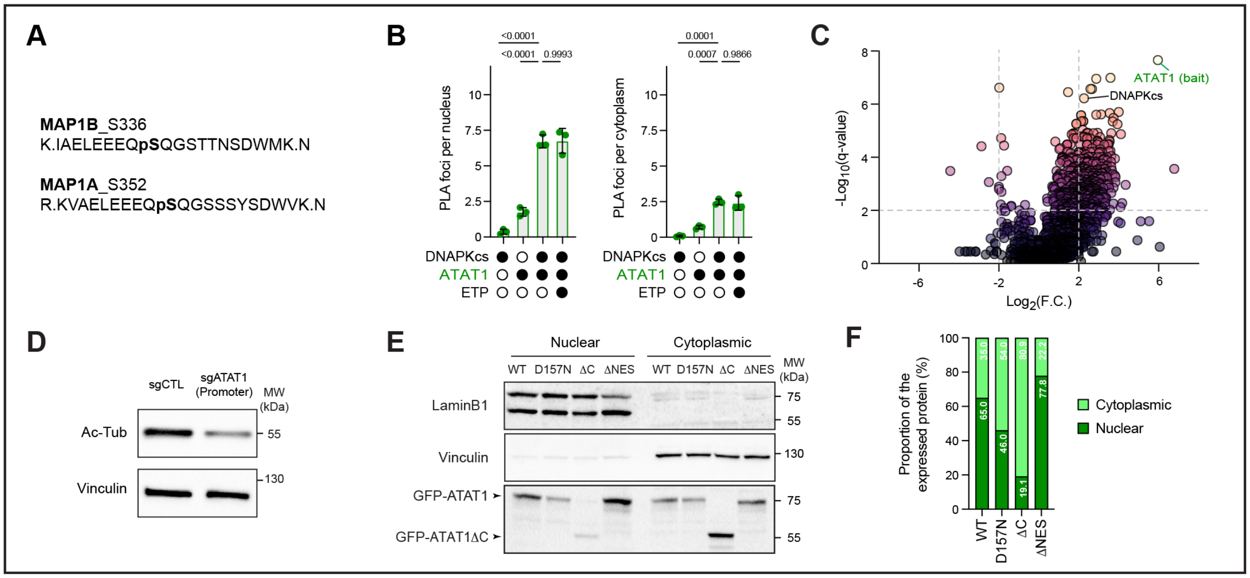


**Supplementary Figure 3. Considerations and controls for protein-protein interactions and ATAT1 mutants; related to Figure 3.** (A) Mass spectrometry identification of MAP1B among ATM- and ATR-phosphorylated proteins. (B) Quantification of nuclear and cytoplasmic DNA-PKcs–ATAT1 PLA foci. (C) Volcano plot of GFP-ATAT1 immunoprecipitation–mass spectrometry (IP-MS) enrichment relative to GFP control. Dashed lines indicate q = 0.01 and twofold enrichment thresholds. (D) Immunoblot analysis of Ac-Tub in ATAT1 knockout cells generated using a promoter-targeting guide RNA. (E, F) Nuclear and cytoplasmic fractionation immunoblots (E) and quantification (F) of GFP-ATAT1 mutant localization, including ΔC and ΔNES constructs. (A-F) Experiments were performed in HEK293 (A), U2OS (B and D–F) or HEK293T (C) cells. GFP-ATAT1 fractionation signals were normalized to LMNB1 (nuclear) or vinculin (cytoplasmic) and expressed as percentages of total normalized signal; PLA data are mean ± SD; n = 3 (B, C) or 2 (E,F) biological replicates; Two-way ANOVA with Tukey's multiple comparisons test (B); IP-MS comparisons (C) were analyzed by unpaired t-tests with two-stage Benjamini–Krieger–Yekutieli false discovery rate correction.

**Supplementary Table 1. siRNA and sgRNA sequences used in this study.**

| Target | Type | Sense Sequence (5’ à 3’) |
|---|---|---|
| CTL | siRNA | Silencer™ Select Negative Control No. 1 siRNA (Cat# 4390843, Ambion) |
| ATAT1 | siRNA | GGCUCAUAAUGAGGUAGAAtt |
| ATAT1 | sgRNA | CGACCGCGCACAATGGGCCA |
| VASH1 | siRNA | GAACAGCCGCAGUGAAAGAtt |
| VASH2 | siRNA | UGAGAAUGAAGAUCCUGAAtt |
| CCTα | siRNA | GAAUUGUGCGGGAUUAUGAtt |
| CCTα 3’UTR | siRNA | GCUGAUAGCUAGAGCCAAAUU |
| CCTβ | siRNA | CAGACACUGUCGCUACGUAtt |
| DGAT1 | siRNA | AGACACUUCUACAAGCCCAtt |
| DGAT2 | siRNA | AGGUCACAGUGGGUCCGAAtt |
| BRCA1 | siRNA | CAACAUGCCCACAGAUCAA<br>CCAAAGCGAGCAAGAGAAU<br>UGAUAAAGCUCCAGCAGGA<br>GAAGGAGCUUUCAUCAUUC |
| XRCC4 | siRNA | GUCUUGAUGUCACUGAUAUtt |
| SUN1 | siRNA | GUGUUGAACUGGGCAAGCAtt |
| KIF5B | siRNA | CCAAUGCUGUGACUUUCGAtt |

**Supplementary Table 2. Plasmids used in this study.**

| Plasmid | Source |
|---|---|
| psPAX-2 | Addgene cat# 12260 |
| pMD2.G | Addgene cat# 12259 |
| pLentiCRISPR v2 | Addgene cat# 52961 |
| pCBASceI | Addgene cat# 26477 |
| pCAGGS-NZEGFP | Gift from Jeremy Stark |
| pCAGGS-BSKX | Gift from Jeremy Stark |
| pEF5B-FRT-GFP-αTAT1 | Addgene cat# 27099 |
| pEF5B-FRT-GFP-αTAT1[D157N] | Addgene cat# 27100 |
| pEF5B-FRT-GFP | This study |
| pEF5B-FRT-GFP-αTAT1[ΔC] | This study |
| pEF5B-FRT-GFP-αTAT1[ΔNES] | This study |
| pcDNA3.1(+)-EGFP | This study |
| pcDNA3.1(+)-EGFP-CCTα (WT) | This study |
| pcDNA3.1(+)-EGFP-CCTα S315A, S319A, S321A, S322A, S323A (pn) | This study |
| pcDNA3.1(+)-EGFP-CCTα S315D, S319D, S321D, S322D, S323D (pm) | This study |
| pcDNA3.1(+)-EGFP-CCTα K122R (cd) | This study |

**Supplementary Table 3. Primers used in this study.**

| Primer | Sequence (5’ à 3’) | Application |
|---|---|---|
| GFP EV FWD | TAACAACTTTCTTGTACAAAGTG | Site-directed mutagenesis GFP empty vector ATAT1 |
| GFP REV | CTCATCAACTTTTTTGTACAAAC | Site-directed mutagenesis GFP empty vector ATAT1 |
| ΔC FWD | TAACAACTTTCTTGTACAAAGTG | Site-directed mutagenesis ΔC GFP-ATAT1 |
| ΔC REV | CCGATGTTGATGGGC | Site-directed mutagenesis ΔC GFP-ATAT1 |
| ΔNES FWD | TGCCCCCCACACCCTAC | Site-directed mutagenesis ΔNES GFP-ATAT1 |
| ΔNES REV | GGGACCTCGTTCTGGTGAG | Site-directed mutagenesis ΔNES GFP-ATAT1 |
| EGFP EV FWD | GGAGGTTCTACCGGTATGGCTGAGC | Site-directed mutagenesis EGFP empty vector CCTα |
| EGFP EV REV | CCCTCTAGACTCGAGTCAGTTAAC | Site-directed mutagenesis EGFP empty vector CCTα |
| S315D, S319D FWD | AGCAGGACCCCAGCAGCAG | Site-directed mutagenesis phosphomimetic CCTα-EGFP |
| S315D, S319D REV | TCGGGTCGATGGCCTGCAGC | Site-directed mutagenesis phosphomimetic CCTα-EGFP |
| S321D, S322D, S323D FWD | GGACCCCGACGATGACCCTACTCGCGAGCGCTC | Site-directed mutagenesis phosphomimetic CCTα-EGFP |
| S321D, S322D, S323D REV | TGCTTCGGGTCGATGGCCTGCA | Site-directed mutagenesis phosphomimetic CCTα-EGFP |
| K122R FWD | ACACAACTTCAGAGGCTTCACGGTGATG | Site-directed mutagenesis catalytic dead CCTα-EGFP |

| | | |
|---|---|---|
| K122R REV | GTGAGCTCATCACTGCAAACT | Site-directed mutagenesis catalytic dead CCTα-EGFP |
| VASH2 FWD | CACGTTGTGCTGGGGATTTAC | RT-qPCR |
| VASH2 REV | GAGTCCGAAAAGTCAATGGCT | RT-qPCR |
| CCTβ FWD | CCATGAAAAACTGACCATTGCTC | RT-qPCR |
| CCTβ REV | GCATAAGGGCTCTTGCATGAC | RT-qPCR |
| DGAT1 FWD | TATTGCGGCCAATGTCTTTGC | RT-qPCR |
| DGAT1 REV | CACTGGAGTGATAGACTCAACCA | RT-qPCR |
| DGAT2 FWD | ATTGCTGGCTCATCGCTGT | RT-qPCR |
| DGAT2 REV | GGGAAAGTAGTCTCGAAAGTAGC | RT-qPCR |
| ACTB FWD | CATGTACGTTGCTATCCAGGC | RT-qPCR |
| ACTB REV | CTCCTTAATGTCACGCACGAT | RT-qPCR |
| GAPDH FWD | GGAGCGAGATCCCTCCAAAAT | RT-qPCR |
| GAPDH REV | GGCTGTTGTCATACTTCTCATGG | RT-qPCR |

**Supplementary Table 4. Antibodies used in this study.**

| Antibody | Company | Catalog # and Ref ID | Application |
|---|---|---|---|
| C6ORF134 (ATAT1) antibody | Proteintech | 28828-1-AP AB_2881219 | IF, WB |
| VASH1 Polyclonal antibody | Thermo Fisher Scientific | PA5-96518 AB_2808320 | IF, WB |
| Alpha Tubulin Antibody (Rb) | Proteintech | 11224-1-AP AB_2210206 | IF, WB |
| Mouse Anti-Tubulin, alpha Monoclonal Antibody, Unconjugated, Clone DM1A | Abcam | Ab7291 AB_2241126 | IF, WB |
| Acetyl-α-Tubulin (Lys40) (D20G3) XP® Rabbit mAb | Cell Signaling Technology | 53355 AB_10544694 | WB |
| Rabbit Anti-tubulin, Detyrosinated | Millipore | AB3201 AB_177350 | WB |
| anti-Polyglutamylation Modification mAb (GT335) | AdipoGen | AG-20B-0020 AB_2490210 | WB |
| MAP1B Polyclonal Antibody | Thermo Fisher Scientific | PA5-78052 AB_2736113 | IF |
| GFP tag antibody | Proteintech | 50430-2-AP AB_11042881 | WB, IP |
| Mouse Anti-DNA PKcs Monoclonal Antibody, | Abcam | Ab44815 AB_731982 | IF, WB |

| | | | |
|---|---|---|---|
| Unconjugated, Clone 39131 | | | |
| HA-Tag (C29F4) Rabbit Monoclonal Antibody, | Cell Signaling Technology | 3724 AB_1549585 | WB |
| Vinculin Antibody | Cell Signaling Technology | 4650 AB_10559207 | WB |
| Anti-LaminB1 antibody- Nuclear Envelope Marker | Abcam | Ab16048 AB_443298 | IF |
| LaminB1 antibody | Proteintech | 66095-1-Ig AB_11232208 | IF, WB |
| Mouse Anti-H2A.X, phosphor (Ser139) Monoclonal Antibody, Unconjugated, Clone 9F3 | Abcam | Ab26350 AB_470861 | IF, WB |
| Anti-RPA2 (phospho S4 + S8) antibody | Abcam | AB264116 | IF, WB |
| Rabbit Anti-RPA32/RPA2 Monoclonal Antibody, clone EPR2877Y | Abcam | AB76420 AB_1524336 | WB |
| PCYT1A antibody [EPR3940(2)] | Abcam | AB109263 AB_10859965 | IF, WB |
| PCYT1A/B_pS315+pS319 | Kinexus | ab-pn546 | WB |
| Phospho-Chk1 (Ser345) Antibody | Cell Signaling Technology | 2341S AB_330023 | WB |
| Human Phospho-Chk2 (T68) Antibody | R&D Systems | AF1626 AB_2291844 | WB |
| DNA PK (phospho 2056) antibody [EPR5670] | Abcam | AB124918 AB_11001004 | WB |
| Anti-Chk1 Antibody | Cell Signaling Technology | 2347 AB_330081 | WB |
| Chk2 (A-12) | Santa Cruz | Sc-5278 AB_627259 | WB |
| Goat anti-Mouse IgG (H+L) Highly Cross-Adsorbed Secondary Antibody, Alexa Fluor™ 488 | Thermo Fisher Scientific | A-11001 AB_2534069 | IF |
| Goat anti-Rabbit IgG (H+L) Highly Cross-Adsorbed Secondary Antibody, Alexa Fluor™ 488 | Thermo Fisher Scientific | A-11008 AB_143165 | IF |
| Goat anti-Mouse IgG (H+L) Cross-Adsorbed Secondary Antibody, Alexa Fluor™ 568 | Thermo Fisher Scientific | A-11004 AB_2534072 | IF |
| Goat anti-Rabbit IgG (H+L) Cross-Adsorbed Secondary Antibody, Alexa Fluor™ 568 | Thermo Fisher Scientific | A-11011 AB_143157 | IF |

| | | | |
|---|---|---|---|
| Goat anti-Mouse IgG (H+L) Cross-Adsorbed Secondary Antibody, Alexa Fluor™ 647 | Thermo Fisher Scientific | A-21235<br>AB_2535804 | IF |
| Goat anti-Rabbit IgG (H+L) Cross-Adsorbed Secondary Antibody, Alexa Fluor™ 647 | Thermo Fisher Scientific | A-21244<br>AB_2535812 | IF |
| Goat anti-Mouse IgG (H+L) Cross-Adsorbed Secondary Antibody, Alexa Fluor™ 405 | Thermo Fisher Scientific | A-31553<br>AB_221604 | IF |
